# Orexinergic Input to the Supramammillary Region Links Threat Salience to Approach Behavior and Dopamine Release

**DOI:** 10.64898/2026.07.18.739326

**Authors:** Yosuke Arima, John Gibons, Jorge Mendoza, Beyonce Getachew, Sarah T. Johnson, Zengyou Ye, Satoshi Ikemoto

## Abstract

Orexin (hypocretin) neurons (OXN) broadcast arousal-related signals across the forebrain, yet how their inputs shape motivated behavior remains poorly understood. The supramammillary region (SuM), a posterior hypothalamic structure implicated in arousal, active coping and reward-seeking, receives dense orexin innervation. Here, we define how orexinergic terminals in the SuM (OXN*_SuM_*) integrate threat salience with approach motivational processes. Using viral tracing and RNAscope, we show that SuM neurons express both orexin receptor subtypes, with type 2 receptors predominating. Fiber photometry recordings from OXN*_SuM_* terminals reveal strong activation by footshock and conditioned threat cues, modest responses to salient sensory stimuli, and pronounced suppression during water consumption. Optogenetic stimulation of OXN*_SuM_* terminals reinforces instrumental responding, biases real-time place preference, and elicits rapid, stimulation-locked dopamine release in the nucleus accumbens. These findings identify an orexin-to-SuM pathway that is engaged by uncertainty and threat yet promotes approach behavior through coupling to mesolimbic dopamine, revealing a mechanism linking hypothalamic arousal signals to adaptive motivational responses.

**SIGNIFICANCE STATEMENT:** The supramammillary region (SuM) has emerged as a critical node linking arousal, threat processing, and motivated behavior, yet the sources and functions of its modulatory inputs remain unclear. Here, we identify orexinergic projections to the SuM as a circuit mechanism that couples threat-related salience with approach behavior and nucleus accumbens dopamine signaling. We show that orexin terminals in the SuM are activated by stressful and uncertain stimuli but suppressed during consummatory behavior, and that selective activation of these terminals is positively reinforcing and drives rapid dopamine release. These findings reveal a pathway through which orexin neurons promote active coping and reward-seeking, advancing our understanding of how hypothalamic neuromodulation shapes mesolimbic function and adaptive behavioral responses.

## INTRODUCTION

To survive environmental challenges, animals must rapidly generate adaptive responses, choosing between passive coping (e.g., freezing) or active engagement (e.g., escaping threats or exploring novel environments). While the underlying neural circuits remain poorly understood, the supramammillary region (SuM), a posterior hypothalamic structure, has emerged as a key node promoting active interaction with the environment (Kesner et al., 2022).

SuM neurons are strongly activated by novelty and stressors, including unfamiliar environments, forced swimming, and footshock, but are inhibited during food or water consumption (Pan and McNaughton, 2004; Kesner et al., 2021; Escobedo et al., 2024; Zhang et al., 2026). Despite activation by aversive events, direct excitation of SuM neurons does not increase passive coping or anxiety; instead, it promotes active environmental interaction (e.g., locomotion, digging, jumping) (Farrell et al., 2021; Escobedo et al., 2024). Rodents readily perform instrumental responding for SuM stimulation, which also activates dopaminergic projections to the nucleus accumbens (NAc) (Ikemoto et al., 2004; Ikemoto, 2005; Ikemoto et al., 2006; Kesner et al., 2021). Conversely, inhibiting SuM neurons disrupts sucrose seeking without reducing actual consumption (Kesner et al., 2021), confirming that the SuM supports active, motivated action rather than consummatory behavior.

Hypothalamic orexin (hypocretin) neurons (OXN) project densely to the SuM, representing a potential source of excitatory drive during environmental challenges (Peyron et al., 1998; Nambu et al., 1999). OXN are strongly activated by acute stressors (Ida et al., 2000; Zhu et al., 2002; Reyes et al., 2003; Chen et al., 2014; Gonzalez et al., 2016). Central administration of orexin recruits physiological stress pathways, triggering autonomic arousal and the release of corticotropin-releasing factor, ACTH, and corticosterone (Ida et al., 2000; Kuru et al., 2000; Samson et al., 2002; Sakamoto et al., 2004; Samson et al., 2007)—responses that are blocked by orexin receptor antagonists (Chang et al., 2007; Samson et al., 2007; Yun et al., 2017).

While orexin signaling can sometimes promote passive or anxious responses like grooming and avoidance (Ida et al., 1999; España et al., 2002; Suzuki et al., 2005), substantial evidence links OXN to active coping and motivation. Chronic stress-induced passive behaviors are reversed by activating OXN (Ito et al., 2008; Ito et al., 2009; Kim et al., 2023). Furthermore, OXN respond to motivationally salient cues, including novel objects, food, or drugs of abuse (Harris et al., 2005; Ennaceur, 2010; Gonzalez et al., 2016; Liao et al., 2024). Orexin signaling is required for drug-seeking behaviors, effort-based food-seeking, and drug-evoked dopamine release in the NAc (Harris et al., 2005; España et al., 2010; Gonzalez et al., 2016; Dong et al., 2026). Consequently, the behavioral outcomes of orexin activation likely depend on the specific downstream circuits engaged.

Given their dense innervation and functional overlap, we hypothesized that orexinergic inputs to the SuM (OXN*_SuM_*) are recruited by stressful or uncertain stimuli to promote active environmental engagement via mesolimbic dopamine pathways. Using anatomical, molecular, physiological, and behavioral approaches, we first confirmed synaptic contacts of orexinergic terminals in the SuM and characterized local orexin receptor types 1 and 2 (OX1R and OX2R, respectively) expression. Next, we recorded OXN*_SuM_* terminal activity during footshocks, conditioned threats, operant behavior, and reward consumption. Finally, we tested whether optogenetic stimulation of OXN*_SuM_* terminals supports reinforcement and evokes dopamine release in the NAc. Together, these experiments identify an orexin-to-SuM pathway that integrates threat salience with motivational processes to drive active, dopamine-dependent behavior.

## MATERIALS AND METHODS

### Animals

Adult male and female C57BL/6J mice (Jackson Laboratory) and orexin-Cre mice were used. Orexin-Cre mice (Matsuki et al., 2009) were maintained by crossing with C57BL/6J mice at the NIDA Transgenic Breeding Facility. Mice were 2–4 months old (25–35 g) at the time of surgery. Animals were group-housed in a temperature- and humidity-controlled vivarium (70–74 °F; 35–55% humidity) on a reversed 12:12 h light–dark cycle (lights off 07:00). Food and water were available ad libitum unless otherwise specified. All procedures were approved by the NIDA IRP Animal Care and Use Committee.

### Viral Vectors

The following AAV constructs were used:

- AAV1-hSyn-FLEX-mGFP-2A-Synaptophysin-mRuby (anterograde terminal labeling)
- pAAV9-Syn-FLEX-GCaMP6s-WPRE-V40 (GCaMP expression in orexin terminals)
- pAAV5-Syn-dLight1.3b (dopamine sensor expression in NAc)
- pAAV5-Syn-FLEX-CrimsonR-tdTomato (red-shifted optogenetic stimulation)
- AAV9-Ef1a-DIO-ChR2-EYFP, AAV9-Ef1a-DIO-ChR2-EYFP, and AAV9-Ef1a-DIO eNpHR 3.0-EYFP (optogenetic activation, control, and inhibition)

All vectors were used at titers of approximately 1×10^1^³ gc/mL.

### Stereotaxic Surgeries

Mice were anesthetized with isoflurane (1–2%) and secured in a stereotaxic frame. Viral vectors were delivered through a beveled 34-gauge needle at 50 nL/min using a Micro4 pump (WPI). Injection volumes were 100–150 nL. The needle was left in place for 10 min before withdrawal. Following viral injection, optical fibers (200 µm core, NA 0.37) were implanted 0.2 mm above the target region and secured with Geristore dental cement.

Precise stereotaxic coordinates were employed for targeting specific brain regions during viral injections and fiber implantations. All measurements are given in millimeters relative to bregma and are listed in the order of anteroposterior, mediolateral, and dorsoventral axes:

- NAc: +1.1, +0.8, -4.5
- HOF: -1.7, +1.0, -4.2
- SuM: -2.8, +0.5, -5.0

Injections and fiber placements were unilateral for ChR2 and EYFP groups; NpHR mice received bilateral injections into the hypothalamic orexin field (HOF) and bilateral fibers above SuM to ensure effective inhibition.

The mice were housed undisturbed for 4–5 weeks before behavioral or photometry testing after AAV injections.

### Histology

Mice were deeply anesthetized and perfused with PBS followed by 10% formalin. Brains were post-fixed overnight, cryoprotected in 20% sucrose, frozen, and sectioned coronally at 40 µm. Sections were mounted with ProLong Diamond and imaged using a Keyence BZ-X710 microscope to verify viral expression and fiber placement.

### Orexin Immunohistochemistry

Sections were incubated with rabbit anti-orexin-A (1:2000; Phoenix Pharmaceuticals) followed by donkey anti-rabbit Cy3 (1:1000; Jackson ImmunoResearch). Fluorescence images were acquired on a Keyence microscope.

### Orexin Neuron Terminal Tracing

Orexin-Cre mice received 100 nL of AAV1-FLEX-mGFP-Synaptophysin-mRuby into the HOF. Brains were processed 4–5 weeks later. Confocal images were obtained with an Olympus FV1000 microscope.

### RNAscope In Situ Hybridization

Wild-type C57BL/6J mice were perfused with 10% formaldehyde and post-fixed for 2 h. Brains were cryoprotected in 20% then 30% sucrose, sectioned at 14 µm, and processed using RNAscope (ACD Bio) probes targeting OX1R and OX2R mRNAs. DapB served as a negative control. Images were collected using a Zeiss LSM 880 confocal microscope.

Quantification was performed using FIJI and QuPath. DAPI-labeled nuclei were detected first, followed by automated puncta detection using fixed thresholds. Cells were classified as receptor-positive if they contained ≥4 puncta or ≥0.4 µm^2^ RNA signal. Counts were made within 900 µm × 600 µm SuM ROIs at anterior and posterior levels (n = 4 mice).

### Fiber Photometry

Dual-wavelength fiber photometry (Doric Lenses) was used to measure the sensor activity. Excitation light consisted of 465 nm sinusoid (modulated at 530 Hz) to excite GCaMP or dLight, and 405 nm sinusoid (modulated at 208 Hz) as an isosbestic control. Both emission signals were collected through a single green emission window (centered at ∼525 nm) and decoupled via frequency demodulation. Signals were sampled at 1200 Hz and synchronized with behavioral events.

Using a customized Python script, the 405 nm control signal was linearly fitted to the 465 nm signal to account for photobleaching and motion artifacts. ΔF/F was calculated as: ΔF/F = (F_(465nm) − F_(fitted control)) / F_(fitted control). Finally, signals were aligned to TTL markers for behavioral events, and subsequent z-score normalization was performed using a pre-event baseline period.

### Water-Seeking Task

The mice were placed on moderate water restriction (15 min free water access per day). Training occurred in two stages: first, the mice underwent a Magazine training in which nose pokes at the reward port (reNP) produced a 5-µL water reward and click sound. Sessions lasted 60 min for five days. Then, they were trained on and subsequently tested with a Two-tone discrimination task. An operant nose poke (opNP) extinguished the house light and triggered a 2 s CS+ (15 kHz) or CS− (3 kHz). CS+ allowed a reNP to deliver 5 µL of water, whereas CS− resulted in no water. Trials resumed when the house light turned on (5 s after CS+/reward or 10 s after CS− offset). Sessions lasted 60 min across eight days. Photometry data obtained on day 8 were used for analysis.

### Salient Sensory-Stimulus Tests

The mice returned to ad libitum water access before sensory tests. In a dark chamber, they received 1-s LED light (55 lux) and 1-s white noise (85 dB) over two sessions. Each stimulus was presented 70 times on a variable interval (45 s). One mouse’s white-noise data were lost due to equipment failure.

### Fear Conditioning

Lastly, the mice underwent two fear conditioning procedures. In a threat acquisition session, the mice were presented with a 6-s cue (light off + tone) was followed 5 s after cue onset by a 1-s, 0.45-mA footshock. Mice received 11 trials per session (variable interval 60 s) for three sessions. In a Shock-omission test, the same cue was followed by shock on 50% of trials (5–7 shock and no-shock trials per mouse).

### Optogenetic Intracranial Self-Stimulation (ICSS)

ICSS was conducted in standard operant chambers equipped with two levers and cue lights. During the first two session, an active-lever press triggered 1-s cue light but no photostimulation. In subsequent sessions, an active-lever press triggered a 16-pulse 50 Hz train, 473-nm photostimulation for ChR2 and EYFP groups and 3-s, 575-nm photostimulation for NpHR groups. Each session lasted 30 min and repeated for 5 or 9 times.

### Real-Time Place Preference (RTPP)

Following ICSS, the mice underwent RTPP in a three-compartment chamber. In session 1 (20 min baseline), no photostimulation was delivered. In session 2, photostimulation was delivered when the mice entered and stayed in compartment A, and in session 3, photostimulation was delivered when the mice entered and stayed in compartment B. ChR2 mice received a 16-pulse 50 Hz train delivered once per second; NpHR mice received 3-s 575-nm illumination once per 4 seconds. Position tracking and stimulation control were managed by EthoVision XT.

### dLight Dopamine Photometry

Mice received dLight or EYFP control AAV into the NAc and Chrimson into the HOF, with a fiber placed above the SuM. After 4–5 weeks, mice were exposed to 15 photostimulation events (50 Hz, 16-pulse train, 638 nm) on a variable interval (45 s) while dLight signals were recorded.

### Experimental design and statistical analyses

Table 1 provides a detailed description of the experimental design and statistical analyses conducted. The results of these analyses are succinctly summarized in the text and figure legends, while Supplementary Table 1 contains extensive results of all analyses conducted.

**Table 1.** Experimental Design and Statistical Analyses (SPSS v31.0.1.0 or GraphPad Prism v.11.02)

| Experiments | DV | Test | Factor Name | N |
| --- | --- | --- | --- | --- |
| RNAscope | Labeled cell count | One-way within-subjects ANOVA | W: Receptor type (OX1R, OX2R, double) | 4 |
| Water-Seeking Task | GCaMP signal AUC | Two-way mixed ANOVA | B: Context (Rewarded, Non-rewarded)<br>W: Epoch (baseline, opNP, CS, reNP, consumption) | Rewarded: 410;<br>Non-rewarded: 385 from 6 mice |
| Salient Sensory-Stimulus Tests | GCaMP signal AUC | t, paired, two-tailed | Light vs. Baseline | 414 events from 6 mice |
|  | GCaMP signal AUC | t, paired, two-tailed; | White noise vs. Baseline | 338 paired events from 5 mice |
| Fear Conditioning | GCaMP signal AUC | Two-way within-subjects ANOVA | W: Phase (ref, CS, shock)<br>W: Epoch (early, late) | 30 paired events from 6 mice |
|  | GCaMP signal AUC | Two-way within-subjects ANOVA | W: Session (1, 3)<br>W: Epoch (ref, CS, shock) | 66 paired events from 6 mice |
|  | GCaMP signal AUC | Two-way within-subjects ANOVA | W: Shock (present, absent)<br>W: Epoch (ref, CS, shock) | 38 paired events from 6 mice |
|  | Photo-stimulation counts | Two-way mixed ANOVA | B: Group (ChR2, EYFP, NpHR)<br>W: Session (3-7) | 10, ChR2;<br>9, EYFP;<br>6, NpHR |
| ICSS | Lever-presses | Two-way within-subjects ANOVA | W: Lever (active, inactive)<br>W: Session (2-7) | 10 |
|  | Active-lever presses | Three-way within-subjects ANOVA | W: Phase (acquisition, extinction, reinstatement)<br>W: Session block (6-7, 8-9, and 10-11)<br>W: Trial (1, 2) | 10 |
|  | Active-lever presses | Two-way within-subjects ANOVA | W: Lever (active, inactive)<br>W: Session (2-7) | 9 |
|  | Active-lever presses | Two-way within-subjects ANOVA | W: Lever (active, inactive)<br>W: Session (2-7) | 6 |
| RTPP | Time (s) | Two-way mixed ANOVA | B: Group (NpHR, ChR2)<br>W: Session (1, 2, 3) | 6, NpHR;<br>10, ChR2 |
| dLight Dopamine Photometry | dLight signal AUC | Two-way mixed ANOVA | AAV (Chrimson, EYFP)<br>Epoch (before, after) | 5, Chrimson;<br>3, EYFP |

Abbreviations: B, between factor; DV, dependent variable; W, Within factor
Note: GCaMP signal data from all trials of individual mice were pooled and analyzed using the described methods.

## RESULTS

### Orexinergic fibers innervate SuM neurons

Previous studies have reported orexin projections to the SuM (Peyron et al., 1998; Nambu et al., 1999). Consistent with this, immunohistochemical staining revealed prominent orexin-immunoreactive fibers and cell bodies within the hypothalamic orexin field (HOF) (Fig. 1A). Within the SuM, orexin-positive fibers were distributed across the anterior–posterior axis, with denser labeling in midline regions (Fig. 1B).

**Figure 1.**
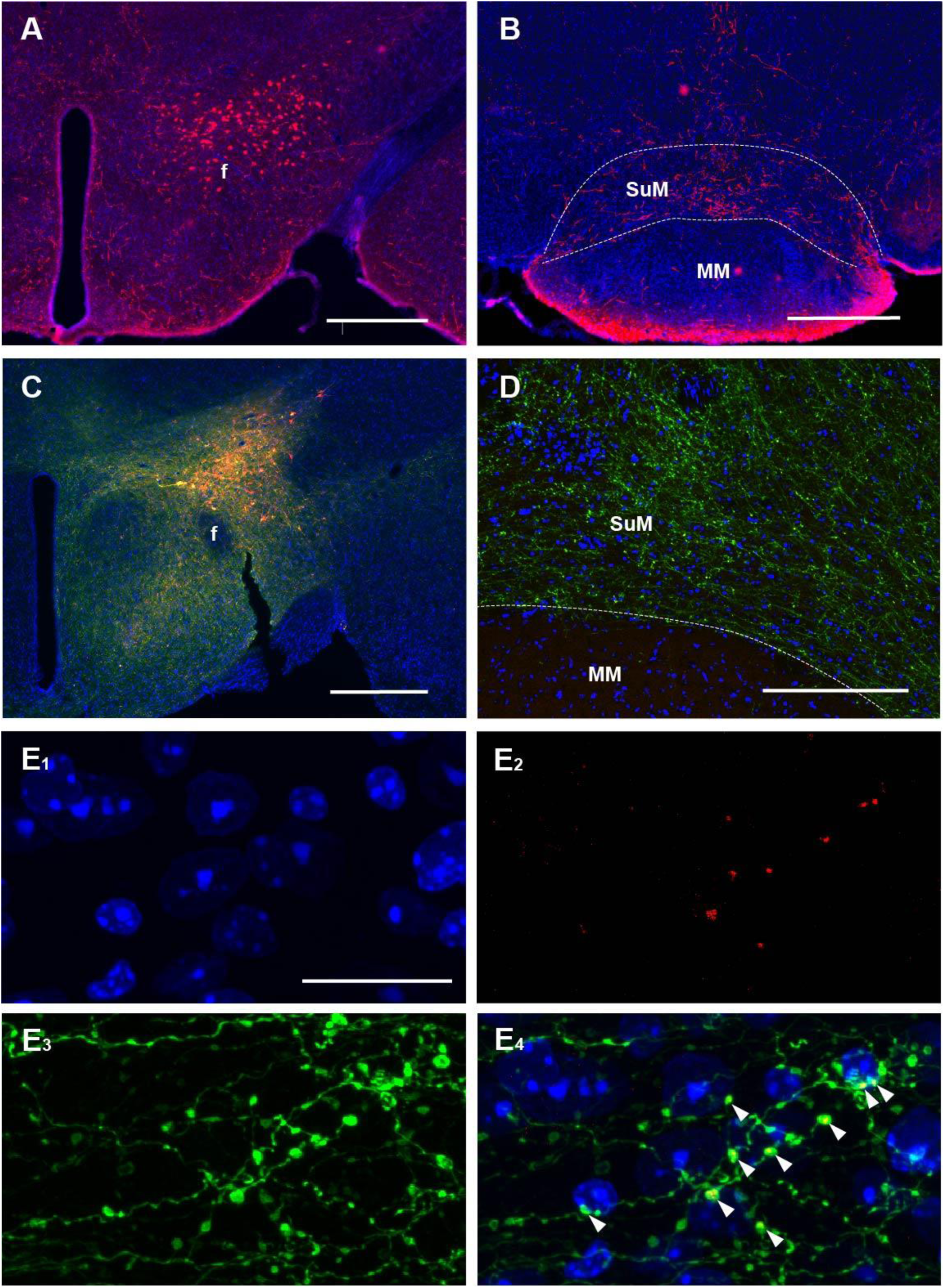
Orexinergic fibers innervate SuM neurons. **(A-B)** Immunohistochemical detection of orexinergic neurons and fibers. Representative photomicrographs of the hypothalamic region show orexin-immunoreactive somata and processes (red) with DAPI-labeled nuclei (blue) at a level of the hypothalamic orexin field (HOF) and supramammillary region (SuM). Scale bar, 500 µm. f, fornix; MM, medial mammillary nucleus. **(C-E)** Viral labeling of orexin neurons and terminals. Injection of AAV1-hSyn-FLEX-mGFP-2A-Synaptophysin-mRuby into the HOF resulted in mGFP-labeled orexinergic somata and axons (green), and synaptophysin-mRuby–labeled presynaptic puncta (red). DAPI labels nuclei (blue). Scale bars: 500 µm (C), 200 µm (D), 20 µm (E). **(E)** Higher-magnification images illustrate mGFP-positive fibers and synaptophysin-mRuby–positive boutons within the SuM. Arrowheads indicate colocalization of synaptophysin with orexinergic axons, including contacts directly apposed to SuM neurons.

To confirm the presence of orexinergic terminals, we used a Cre-dependent viral anterograde tracing strategy. Injection of AAV1-hSyn-FLEX-mGFP-2A-Synaptophysin-mRuby into the HOF of orexin-Cre mice yielded GFP- and mRuby-labeled somata, axons, and terminals in regions dorsal to the fornix (Fig. 1C). GFP-labeled fibers were readily visible in the SuM at low magnification (Fig. 1D), and confocal imaging revealed discrete synaptophysin-mRuby–positive puncta colocalized with GFP-positive fibers (Fig. 1E). These findings indicate that orexin neurons robustly innervate the SuM and form synaptic contacts with SuM neurons.

### SuM neurons express both orexin receptor subtypes, with OX2R predominance

To examine orexin receptor expression in the SuM, we performed RNAscope in situ hybridization for OX1R and OX2R. Transcripts for both receptors were detected throughout the SuM (Fig. 2A). Quantification within a defined 900 µm × 300 µm region across multiple sections revealed an average of 686.8 ± 29.6 DAPI-labeled cells per animal (n = 4). Among these, 20.2% ± 5.4% were OX1R-positive and 56.6% ± 8.1% were OX2R-positive (Fig. 2B).

**Figure 2.**
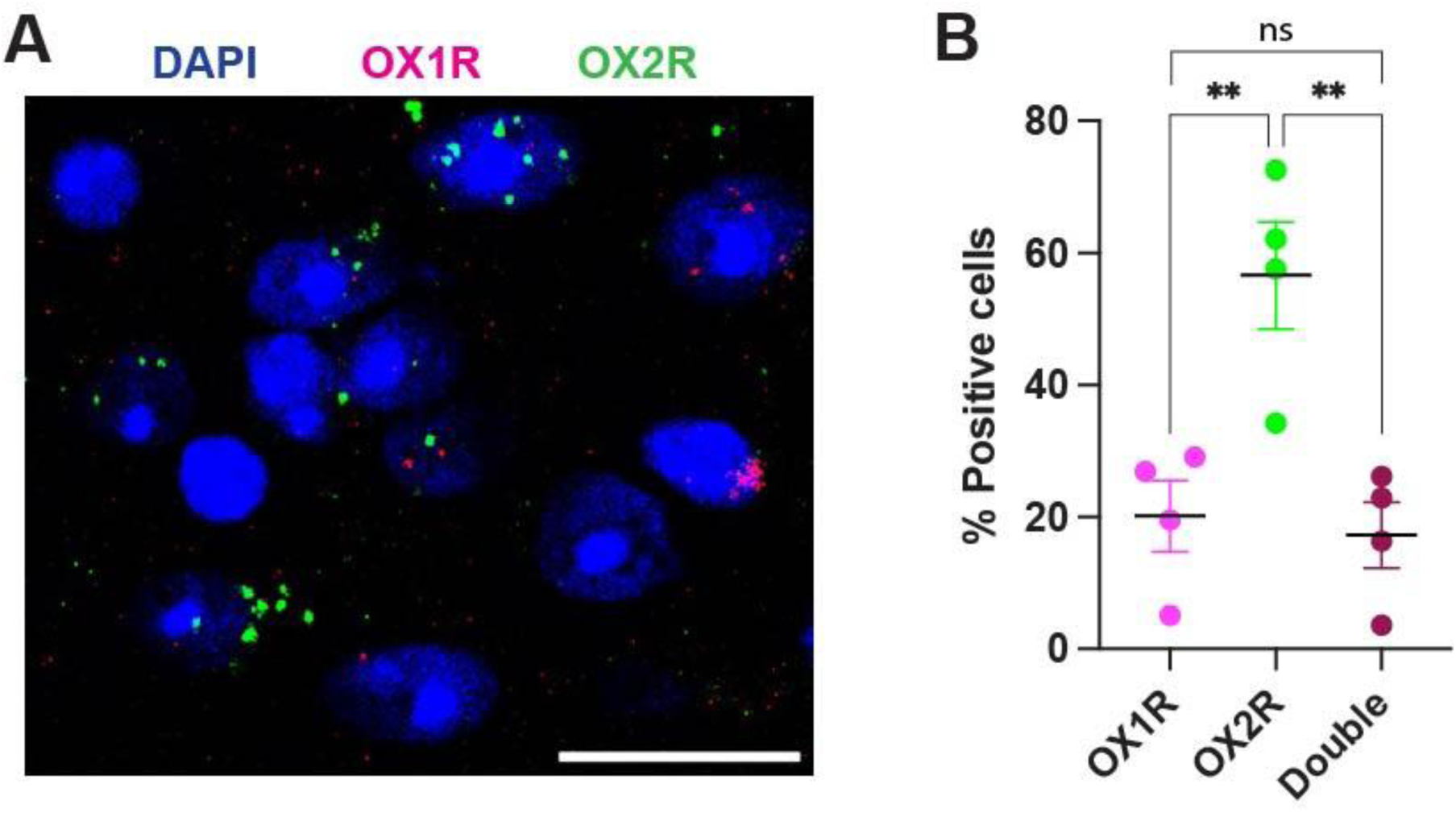
SuM neurons express orexin receptors. **(A)** Representative RNAscope images showing OX1R (magenta) and OX2R (green) mRNA expression in the SuM. Nuclei are labeled with DAPI (blue). Contrast was optimized to improve probe visibility. Scale bar, 500 µm. **(B)** Quantification of receptor expression. Percentages of DAPI-labeled neurons expressing OX1R, OX2R, or both receptors are shown (mean ± SEM; n = 4 mice). **P < 0.01, Tukey’s multiple-comparison test.

A one-way ANOVA revealed a significant receptor-type effect (F(1.2, 3.5) = 115.0, P < 0.001), indicating a clear predominance of OX2R expression (Supplementary Table 1). Notably, 17.3% ± 5.0% of neurons expressed both receptors, and this fraction did not differ significantly from the OX1R-only population, suggesting that most OX1R-positive neurons also coexpress OX2R.

### Orexinergic terminals in the SuM are activated by stressful and salient stimuli and inhibited during consummatory behavior

To characterize the activity of orexinergic terminals in the SuM (OXN*_SuM_*), we used fiber photometry to record GCaMP signals in orexin-Cre mice.

#### Activity during water-seeking behavior

Mice were trained under water restriction to perform an operant water-seeking task (Fig. 3A,B). A 5×2 mixed ANOVA (epoch × context) revealed the following (Fig. 3c,d): Operant nose pokes (opNP) produced a small but reliable increase in OXN*_SuM_* activity. Both conditioned stimuli (CS+ and CS−) generated similar transient increases in activity, indicating that OXN*_SuM_* did not discriminate between reward-predictive and non-predictive cues during this epoch (Fig. 3C,D). Reward nose pokes (reNP) following CS+ produced a significant decrease in activity relative to baseline, whereas reNP following CS− did not, although direct comparison of the reNPs in the two contexts indicated no difference. Post-reNP periods showed suppression in both contexts, with stronger, marked suppression during actual water delivery.

**Figure 3.**
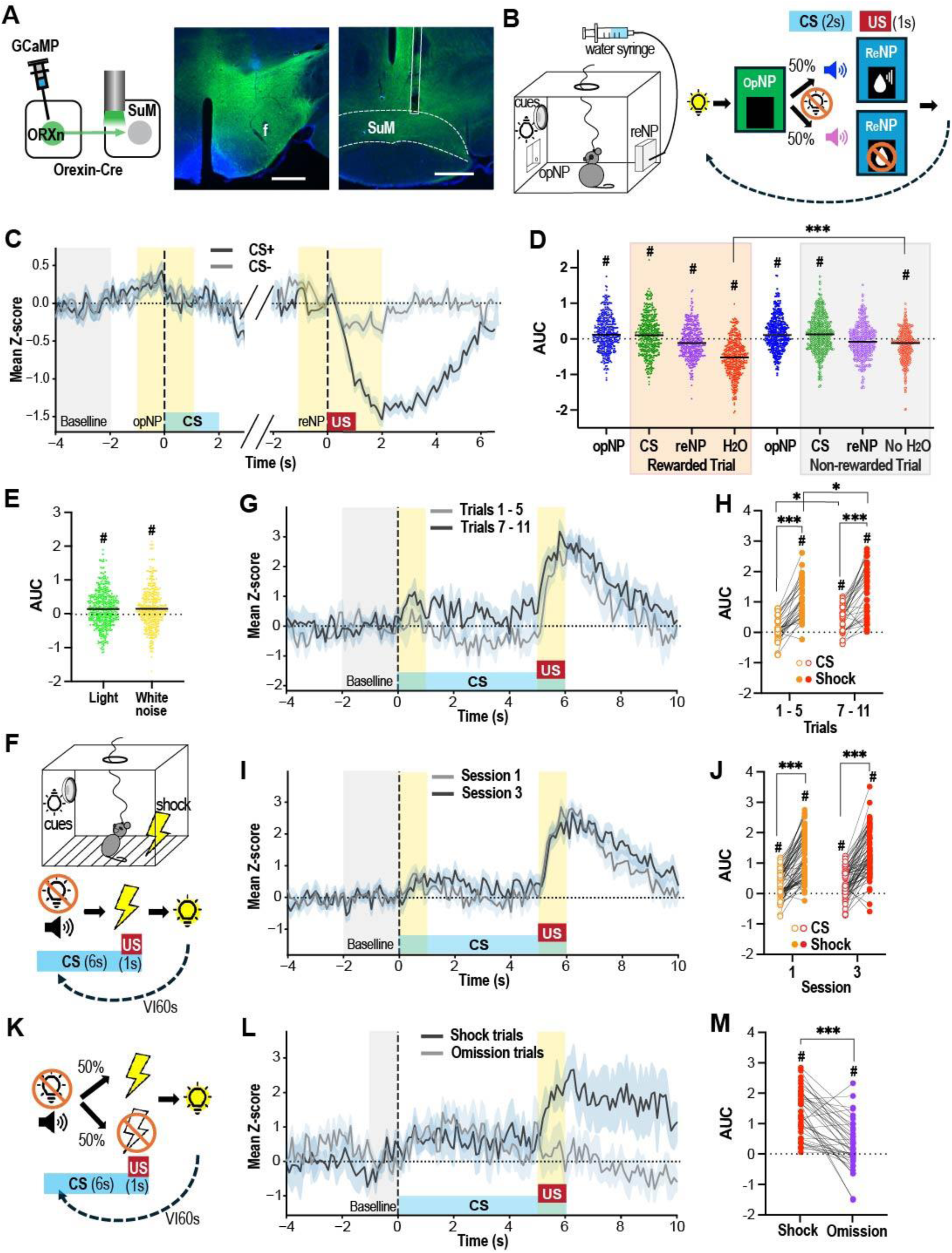
Orexinergic terminals in the SuM are activated by stressful stimuli and suppressed during consummatory behavior. **(A)** Schematic of GCaMP viral injection and optic fiber placement (left), GCaMP expression in the HOF (middle), fiber track above SuM (right). Scale bar, 500 µm. **(B)** Water-seeking task. Left, operant chamber layout including house light, tone speaker, operant nose-poke (opNP) port, and reward nose-poke (reNP) port. Right, task structure: an opNP initiates a trial and triggers conditions stimuli (house-light extinction and 2-s tones). A reNP following CS+ delivers 5 µL of water; reNPs following CS− produce no reward. **(C)** GCaMP fluorescence traces were collected and analyzed in alignment with two distinct behavioral events: the operant nose-poke (opNP) and the reward nose-poke (reNP). To quantify changes in neuronal activity, Z-scores were calculated and expressed as mean ± SEM. The reference window used for establishing baseline activity was set from −4 to −2 seconds prior to opNP. This baseline period provided a standard for comparison against subsequent activity changes. Specific epochs were highlighted with yellow shading to indicate the segments selected for Area Under the Curve (AUC) calculations. These epochs correspond to periods of interest surrounding the behavioral events. For analytic consistency, both the baseline AUC and the post-reNP AUC values were halved prior to statistical analysis. **(D)** AUC values for behavioral epochs. OXN*_SuM_* ativity increased during opNP and during both CS types, and decreased during reNP+ but not reNP− and during water consumption. #P < 0.01 versus baseline AUC; ***P < 0.001; Bonferroni-corrected. **(E)** AUC signal changes to salient sensory stimuli. In a dark chamber, 1-s LED light (55 lux) or 1-s white noise (85 dB) increased GCaMP signals. #P < 0.001 versus baseline (paired t-test). **(F, K)** Schematic of fear-conditioning (F) and shock-omission (K) procedures. The CS consisted of light-off plus a 6-s tone, followed 5 s later by a 1-s, 0.4-mA footshock (except during omission trials). **(G, I, L)** Representative GCaMP traces aligned to CS onset. Early vs. late trials (G), session 1 vs. session 3 comparisons (I), and shock vs. omission responses (l) are shown. Z-score reference windows: −2 to 0 s for (G, I); −1 to 0 s for (L). **(H, J, M)** AUC quantifications. CS responses increased across early to late trials (H). Responses were stable between sessions (J). Shock omission did not decrease GCaMP activity (M). *P < 0.05; ***P < 0.001; #P < 0.05 versus baseline (Bonferroni-corrected).

Overall, OXN*_SuM_* activity increased during operant engagement and early cue processing but decreased sharply during the consummatory phase. Notably, opNP and reNP required essentially the same movements, yet their effects on orexinergic terminal signals were opposite. This suggests that OXN*_SuM_* activity is more closely linked to the animal’s expectations regarding the outcome of their behavior rather than the movements themselves.

#### Activity during salient sensory stimuli

In a dark chamber, the mice were exposed to repeated presentations of bright light or white noise (70 trials, variable interval 45 s). Both stimuli produced small but reliable increases in OXN*_SuM_* activity (Fig. 3E), indicating sensitivity to salient sensory events.

#### Activity during footshock and conditioned threat

The mice then underwent a footshock-based Pavlovian conditioning task (Fig. 3F). During session 1, the CS produced no detectable increase in signals during the first five trials but elicited significant increases in the last five trials, indicating the CS acquiring affective value (Fig. 3G,H). Footshock itself produced a strong increase in activity throughout the session, with slightly larger responses in later trials.

Across sessions 1 and 3, CS-evoked and shock-evoked responses were similar, confirming that conditioning was acquired rapidly (Fig. 3I,J). No session or session × epoch interactions were observed (Supplementary Table 1).

#### Activity during shock-omission trials

During a partial-reinforcement session (50% shock omission; Fig. 3K), OXN*_SuM_* activity increased when the CS was followed by shock omission, although the increase was significantly smaller than when shock was presented (Fig. 3L,M; significant omission × epoch interaction: F(1, 37) = 44.67, P < 0.001; Supplementary Table 1). This suggests that OXN*_SuM_* do not encode negative reward prediction errors.

### Optogenetic activation of OXN*_SuM_* terminals reinforces approach behavior

Given that SuM stimulation is reinforcing (Ikemoto et al., 2004; Ikemoto, 2005; Ikemoto et al., 2006; Kesner et al., 2021), we next tested whether selective activation of OXN*_SuM_* terminals produces similar effects. Orexin-Cre mice received AAV-DIO-ChR2, AAV-DIO-EYFP, or AAV-DIO-eNpHR in the HOF and an optic fiber targeting the SuM (Fig. 4A).

**Figure 4.**
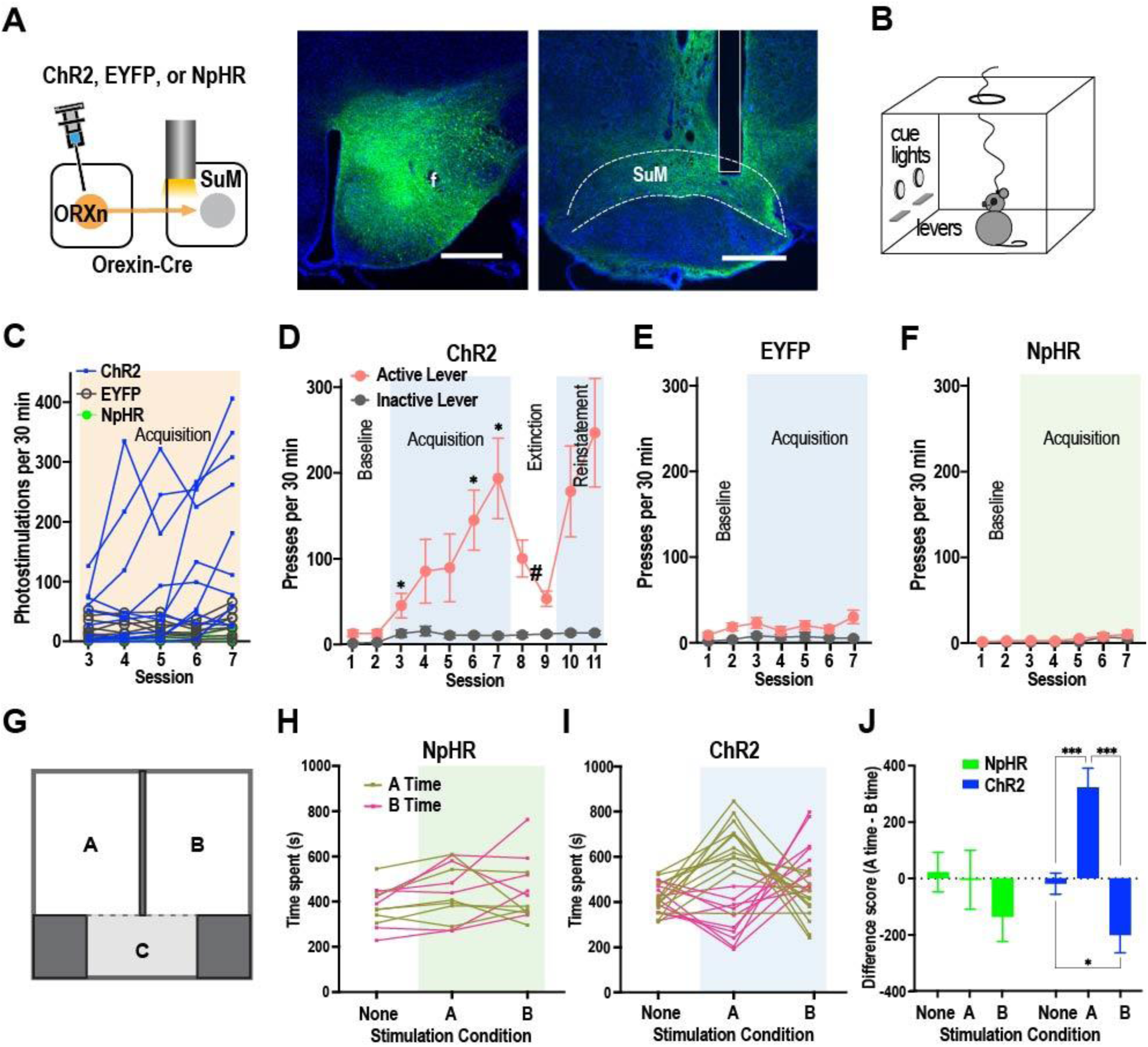
Optogenetic activation of OXN*_SuM_* terminals reinforces approach behavior and induces positive affect. **(A)** Schematic of viral injections and fiber placement (left), EYFP expression in the HOF (middle), and fiber trajectory targeting the SuM (right). **(B)** Operant chamber layout. It included two cue lights and two levers. An active-lever press triggered cue light and photostimulation, while inactive-lever presses resulted in no programmed consequence. **(C)** Photostimulation earned by individual mice during ICSS acquisition (sessions 3–7). ChR2 mice showed progressive increases in earned stimulations, significantly exceeding EYFP and NpHR controls in sessions 6–7 (Ps < 0.005; Bonferroni-corrected). **(D-F)** Active and inactive lever presses (mean ± SEM). ChR2 mice selectively increased active-lever responding across acquisition and reinstatement phases. EYFP controls displayed mild cue-related bias, while NpHR mice showed no lever preference. *Ps < 0.05; #Ps < 0.05 versus session blocks 6-7 and 10-11 (Bonferroni-corrected). **(G)** Place-preference chamber layout. The chamber consists of three compartments between which the mouse was able to move freely. Compartments A and B are used for photostimulation. **(H-J)** Real-time place preference. NpHR mice showed no preference or avoidance during stimulation in either compartment (H). ChR2 mice displayed robust preference for the stimulation-paired compartment (I). Difference scores (A – B) across conditions confirmed that ChR2 mice showed consistent preference and NpHR mice showed none (J). *P = 0.043; ***P < 0.001 (Bonferroni-corrected).

#### Intracranial self-stimulation

ChR2 mice earned significantly more photostimulation events than EYFP or NpHR controls (Fig. 4B,C). A mixed ANOVA revealed a significant group × session interaction (F(8, 88) = 3.76, P < 0.001). During the acquisition phase, ChR2 mice developed a robust active-lever preference (Fig. 4D), with a significant lever × session interaction (F(2.5, 45.5) = 7.17, P < 0.001). EYFP controls showed only a mild, cue-related lever bias, while NpHR mice showed no lever discrimination (Fig. 4E,F). Together, these results indicate that activation—but not inhibition—of OXN*_SuM_* terminals is reinforcing.

#### Real-time place preference

To test whether inhibition of OXN*_SuM_* is aversive, the NpHR and ChR2 mice underwent a real-time place-preference test (Fig. 4G). NpHR mice showed no avoidance or preference for stimulation-paired compartments (Fig. 4H), whereas ChR2 mice showed robust preference for the stimulation-paired compartment across both stimulation conditions (Fig. 4I). A group × condition ANOVA confirmed a significant interaction (F(2, 28) = 6.20, P = 0.006) (Fig. 4J).

Thus, while activation of OXN*_SuM_* produces a positive affective state, inhibition does not result in aversion.

### Optogenetic activation of OXN*_SuM_* terminals evokes time-locked dopamine release in the nucleus accumbens

To determine whether OXN*_SuM_* activation influences mesolimbic dopamine signaling, we performed dLight fiber photometry in the NAc while stimulating OXN terminals in the SuM (Fig. 5A-D).

**Figure 5.**
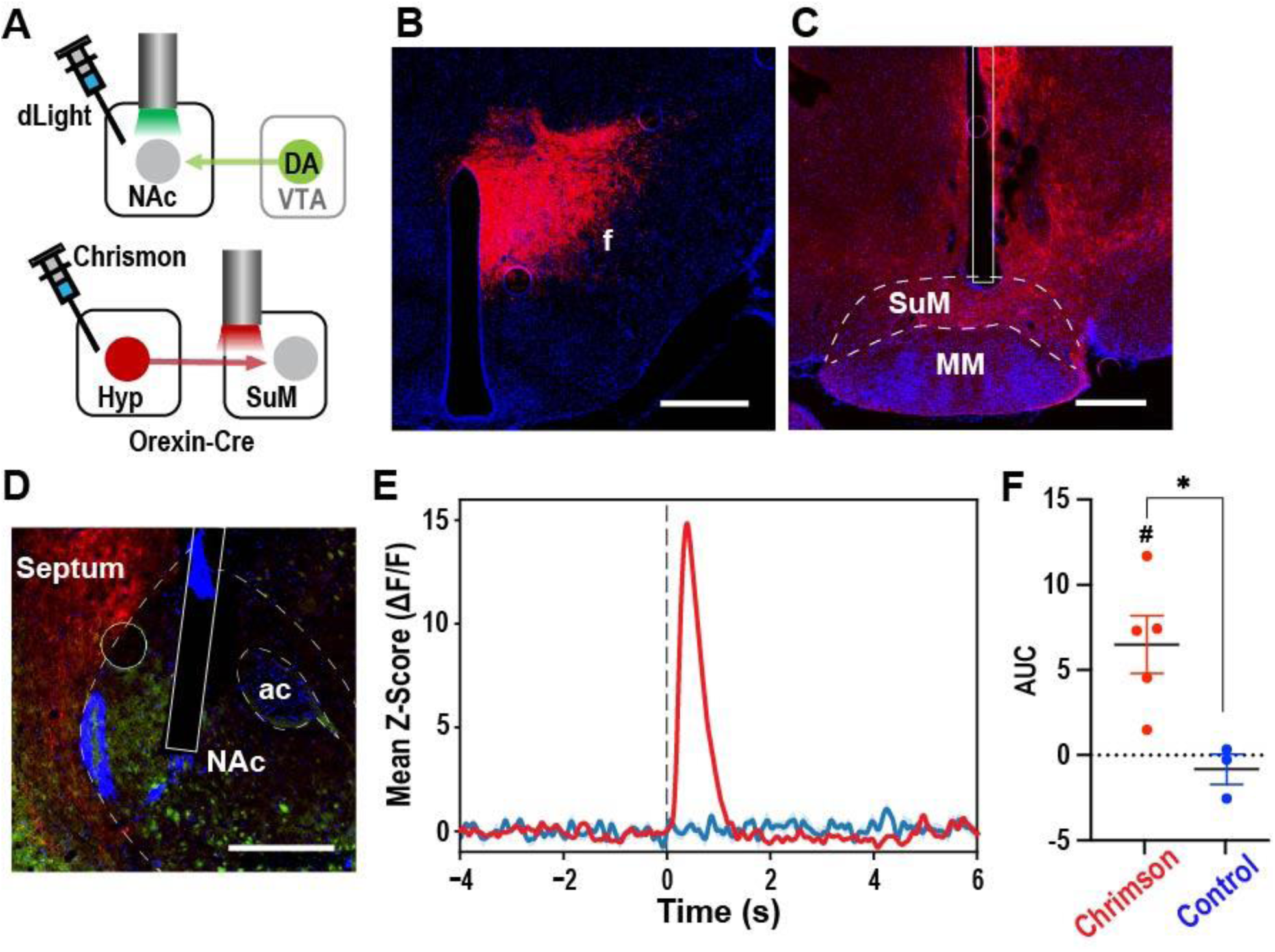
Activation of OXN*_SuM_* terminals evokes stimulation-locked dopamine release in the nucleus accumbens. **(A-D)** Schematic of dLight and Chrimson injections and fiber placements (A). Chrimson-labeled orexin neurons (red) in HOF (B). Fiber targeting the SuM (C). dLight expression (green) in the nucleus accumbens (D). Scale bars, 500 µm. ac, anterior commissure; DA, dopamine; f, fornix; MM, medial mammillary nucleus; VTA, ventral tegmental area. **(E)** Representative z-scored dLight traces. Time 0 s indicates the onset of photostimulation. In Chrimson-expressing mice, SuM photostimulation (16-pulse 50 Hz train) elicited a rapid, time-locked increase in dopamine signal (red). Control mice showed no change (blue). **(F)** Mean AUC (0–2 s) in relation to baseline (−2–0 s). Chrimson mice showed significant stimulation-evoked dopamine release; controls did not. *P = 0.042; #P = 0.004 versus baseline (Bonferroni-corrected). Group sizes: Chrimson, n = 5; controls, n = 3.

Photostimulation (16-pulse train, 50 Hz) produced a robust, time-locked increase in dLight fluorescence in ChR2-expressing mice (Fig. 5E,F) with no detectable change in control mice. A two-way ANOVA revealed a significant epoch × treatment interaction (F(1, 6) = 9.71, P = 0.021; Supplementary Table 1). These data show that OXN*_SuM_* terminals can rapidly drive NAc dopamine release.

## DISCUSSION

Here, we identify a novel functional role for orexinergic projections to the supramammillary region (OXN_SuM_) in linking threat-related environmental activation with positive reinforcement and mesolimbic dopamine signaling. Using a convergent approach spanning anatomy, molecular biology, physiology, and behavior, we demonstrate that orexin neurons densely innervate the SuM, where post-synaptic neurons express both orexin receptor subtypes with a marked predominance of OX_2_R. In vivo fiber photometry revealed a distinct activity pattern in these terminals: they are rapidly recruited by salient or uncertain environmental stimuli, but actively suppressed during consummatory behavior. Finally, selective optogenetic activation of OXN_SuM_ terminals is reinforcing and drives rapid dopamine release in the nucleus accumbens (NAc), establishing a functional hypothalamic-mesolimbic circuit through which orexinergic signaling coordinates active, motivated behavioral states.

### Anatomical and Molecular Organization of the OXN_SuM_ Pathway

Immunohistochemistry and Cre-dependent anterograde tracing showed abundant orexin-immunoreactive fibers and synaptophysin-positive presynaptic terminals in the SuM, indicating strong excitatory input from OXN. RNAscope assays revealed expression of both OX1R and OX2R mRNAs, with OX2R nearly twice as prevalent as OX1R, and substantial overlap between the two receptor populations.

OX_2_R signaling is well characterized for its critical role in arousal, vigilance, and wakefulness (Chieffi et al., 2017), whereas OX_1_R signaling has been heavily implicated in reward processing, motivation, and the modulation of dopaminergic pathways (James et al., 2017). The extensive co-expression of OX1R and OX2R suggests that the SuM may function as an integrative node where vigilance and arousal states are translated into motivational output through downstream dopamine pathways. Future studies employing pathway-specific receptor knockdown or pharmacology will be crucial to dissect how local OX_1_R- and OX_2_R-mediated intracellular cascades differentially contribute to the distinct behavioral components of SuM activation.

### Dynamic Coding of Environmental Salience and Behavioral State

Our fiber photometry recordings revealed that OXN*_SuM_* terminals dynamically encode environmental salience rather than specific hedonic valence. Terminals exhibited robust activation in response to unconditioned stressors, including footshock, bright light, and loud noise. Furthermore, previously neutral stimuli acquired the capacity to activate OXN*_SuM_* terminals after pairing with footshock, indicating that this pathway processes both primary physical threats and learned environmental predictors of danger.

During operant water-seeking behavior, we observed modest increases in OXN*_SuM_* activity during active operant engagement and cue presentation (CS^+^ and CS^-^). Conversely, terminal activity decreased markedly during reward consumption, a pattern that closely parallels general orexin neuron dynamics during ingestive behavior (Dong et al., 2026). This transition suggests that the OXN*_SuM_* pathway is preferentially recruited during active, externally oriented behavioral states—such as exploration, cue evaluation, and threat monitoring—and is actively suppressed during consummatory phases when active environmental engagement is no longer required.

The absence of activity changes during shock omission indicates that OXN *_SuM_* terminals do not encode negative prediction errors. Their functional profile corresponds to real-time encoding of environmental salience, uncertainty, and behavioral mobilization, rather than specific reward or punishment outcomes.

Identifying the inhibitory pathways that silence OXN*_SuM_* activity during eating or drinking (e.g., GABAergic projections from hypothalamic or forebrain circuits) will be important for understanding how seeking behavior is terminated when a goal is achieved.

### Functional Integration into Broad Networks of Active Coping

Optogenetic stimulation of OXN *_SuM_* terminals was found to reinforce instrumental responding and produce strong real-time place preference. This outcome demonstrates a positive affective effect associated with terminal activation. In contrast, optogenetic inhibition did not induce aversion, indicating that tonic orexinergic input is not necessary for maintaining baseline affective state. Therefore, OXN *_SuM_* terminals appear to provide an affectively positive motivational drive that encourages approach behavior when activated.

Stimulation of OXN*_SuM_* terminals evoked rapid, stimulation-locked dopamine release in the NAc. This response occurred within seconds and was absent in controls, demonstrating that orexin input to the SuM can immediately engage the mesolimbic dopamine system. Prior work shows that SuM neurons influence ventral tegmental area (VTA) dopamine neurons through medial septal glutamatergic pathways (Kesner et al., 2021). Given this, it is plausible that activation of orexinergic terminals leads to activation of SuM-MS-VTA pathways, resulting in mesolimbic dopamine release.

These findings align with and extend the broader framework of orexinergic regulation of active coping and motivational engagement. Orexin neurons are known to be recruited by acute stressors, novelty, and motivationally salient cues, and orexin signaling is required for both the reinstatement of drug-seeking behaviors and physical arousal under threat. Our data locate the SuM as a critical downstream node through which these broad orexinergic signals are channeled to generate specific, active behavioral responses.

### Limitations

Several limitations should be considered. Although SuM neurons express OX1R and OX2R mRNAs, receptor protein expression and functional engagement were not examined, leaving the relative contributions of each receptor subtype unresolved. Additionally, orexin neurons release multiple neurotransmitters—including glutamate (Rosin et al., 2003; Torrealba et al., 2003), dynorphin (Chou et al., 2001), and nociceptin/orphanin FQ (Maolood and Meister, 2010)—and it remains unclear which transmitters contribute to threat-evoked or reinforcement-related effects at OXNSuM terminals. Finally, although OXN*_SuM_* terminals were activated by threat and produced reinforcement when stimulated, it is not yet known whether the same terminals mediate both processes or whether distinct subsets support threat versus reward functions. Similarly, SuM neurons engaged by threat may differ from those supporting reinforcement.

### Closing Remarks

In summary, orexinergic input to the SuM is activated by salient and threatening stimuli, suppressed during consummatory behavior, and capable of driving reinforcement and dopamine release when stimulated. These findings identify the OXN–SuM pathway as a mechanism influencing approach-related motivational processes under threats. Determining how orexin signaling interacts with SuM output pathways will be important for clarifying how stress and motivational states shape behavior and may reveal how dysregulation in this circuit contributes to maladaptive responses in chronic stress, depression, or PTSD.

## Acknowledgments

The authors thank Dr. Catherine M. Kotz (University of Minnesota) for generously providing breeding pairs of orexin-Cre mice with the permission from Dr. Takeshi Sakurai (Tsukuba University). The authors also acknowledge the use of the microscopy facilities provided by the NIDA Microscope Core.

The contributions of the NIH authors are considered Works of the United States Government. The findings and conclusions presented in this paper are those of the author(s) and do not necessarily reflect the views of the NIH or the U.S. Department of Health and Human Services.

