## Supplemental Table 1 for "Orexinergic Input to the Supramammillary Region Links Threat Salience to Approach Behavior and Dopamine Release"

Supplementary Table 1. Statistical Results (SPSS v31.0.1.0 or GraphPad Prism v.11.02)

| Figure | DV | Test | Factor Name | Value | P Value | Partial Eta$^2$ |
|---|---|---|---|---|---|---|
| 2B | Labeled cell count | ANOVA, within-subjects design with receptor type; n = 4 | Receptor type | $F_{1.2, 3.5}$ = 115.0 | < 0.001 | |
| 3D | GCaMP signal AUC | ANOVA, mixed design with Context (Rewarded, Non-rewarded), Epoch (baseline, opNP, CS, reNP, compumption); n = 410 and 385 (from 6 mice) for rewarded and non-rewarded, respectively | Epoch (sphericity)<br>Context (between)<br>Epoch (within)<br>Context x Epoch | $\chi^2$ = 138.19<br>$F_{2, 22}$ = 15.79<br>$F_{2.2, 47.8}$ = 36.67<br>$F_{8, 88}$ = 4.74 | < 0.001<br>< 0.001<br>< 0.001<br>< 0.001 | 0.92$^E$<br>0.02<br>0.24<br>0.04 |
| 3E | GCaMP signal AUC | t, paired, two-tailed; 414 events from 6 mice | Light vs. Ref. | $t_{1, 413}$ = 6.06 | < 0.001 | 0.08 |
| | GCaMP signal AUC | t, paired, two-tailed; 338 paired events from 5 mice | White noise vs. Ref | $t_{1, 337}$ = 4.16 | < 0.001 | 0.05 |
| 3H | GCaMP signal AUC | ANOVA, within-subjects design with epoch (ref, CS, shock), phase (early, late); 30 paired events from 6 mice | Epoch (sphericity)<br>Phase (within)<br>Epoch (within)<br>Phase x Epoch | $\chi^2$ = 24.48<br>$F_{1, 29}$ = 11.67<br>$F_{1.3, 36.6}$ = 88.9<br>$F_{2, 58}$ = 4.48 | < 0.01<br>= 0.002<br>< 0.001<br>= 0.016 | 0.63$^E$<br>0.29<br>0.75<br>0.13 |
| 3J | GCaMP signal AUC | ANOVA, within-subjects design with epoch (ref, CS, shock), session (1, 3); 66 paired events from 6 mice | Epoch (sphericity)<br>Session x Epoch (sphericity)<br>Session (within)<br>Epoch (withing)<br>Session x Epoch | $\chi^2$ = 30.53<br>$\chi^2$ = 10.60<br><br>$F_{1, 65}$ = 1.36<br>$F_{1.5, 94.3}$ = 201.28<br>$F_{1.7, 112.79}$ = 0.84 | < 0.01<br>= 0.005<br><br>= 0.247<br>= 0.001<br>= 0.421 | 0.73$^E$<br>0.87$^E$<br><br>0.02<br>0.76<br>0.01 |
| 3M | GCaMP signal AUC | ANOVA, within-subjects design with omission (yes, no), epoch (ref, CS, shock); 38 paired events from 6 mice | Stimulus (within)<br>Epoch (withing)<br>Stimulus x Epoch | $F_{1, 37}$ = 43.26<br>$F_{1, 37}$ = 55.33<br>$F_{1, 37}$ = 44.67 | < 0.001<br>< 0.001<br>< 0.001 | 0.54<br>0.60<br>0.55 |
| 4C | Photo-stimulation counts | ANOVA, mixed design with group (ChR2, EYFP, NpHR), session (5); n = 10, 9, and 6 for ChR2, EYFP, and NpHR, respectively | Session (sphericity)<br>Group (between)<br>Session (within)<br>Group x Session | $\chi^2$ = 39.06<br>$F_{2, 22}$ = 7.99<br>$F_{2.2, 47.8}$ = 4.25<br>$F_{8, 88}$ = 3.76 | < 0.01<br>= 0.002<br>= 0.017<br>< 0.001 | 0.54$^E$<br>0.42<br>0.16<br>0.26 |
| 4D | Lever-presses | ANOVA, within-subjects design with lever (2) and session (5); n = 10 | Lever (within)<br>Session (within)<br>Lever x Session | $F_{1, 18}$ = 11.50<br>$F_{2.5, 45.5}$ = 7.83<br>$F_{2.5, 45.5}$ = 7.17 | = 0.003<br>< 0.001<br>< 0.001 | |
| | Active lever-presses | ANOVA, within-subjects design with phase (acquisition, extinction, and reinstatement) and session (6-7, 8-9, and 10-11); n = 10 | Phase (sphericity)<br>Phase (within)<br>Session (within)<br>Phase x Session | $\chi^2$ = 7.53<br>$F_{1.2, 11.2}$ = 8.48<br>$F_{1, 9}$ = 11.45<br>$F_{2, 18}$ = 8.22 | < 0.023<br>= 0.011<br>= 0.008<br>= 0.003 | 0.62$^E$<br>0.49<br>0.56<br>0.48 |
| 4E | Lever-presses | ANOVA, within-subjects design with lever (2) and session (5); n = 9 | Lever (within)<br>Session (within)<br>Lever x Session | $F_{1, 16}$ = 7.08<br>$F_{3.1, 49.3}$ = 2.48<br>$F_{3.1, 49.3}$ = 2.53 | = 0.017<br>= 0.070<br>= 0.066 | |
| 4F | Lever-presses | ANOVA, within-subjects design with lever (2) and session (5); n = 6 | Lever (within)<br>Session (within)<br>Lever x Session | $F_{1, 10}$ = 1.39<br>$F_{2.2, 21.7}$ = 3.83<br>$F_{2.2, 21.7}$ = 0.29 | = 0.265<br>= 0.035<br>= 0.766 | |
| 4J | Time (s) | ANOVA, within-subjects design with group (2) and stimulation condition (3); n = 6 and 10 for NpHR and ChR2, respectively | Condition (sphericity)<br>Group (between)<br>Condition (within)<br>Group x Condition | $\chi^2$ = 9.35<br>$F_{1, 14}$ = 1.10<br>$F_{1.3, 18.5}$ = 4.06<br>$F_{2, 28}$ = 6.20 | = 0.009<br>= 0.311<br>< 0.001<br>= 0.006 | 0.66$^E$<br>0.07<br>0.50<br>0.31 |
| 5F | dLight signal AUC | ANOVA, mixed design with AAV (Chrimson, EYFP), epoch (before, after); n = 5 and 3 for Chrimson and EYFP, respectively | AAV (between)<br>Epoch (within)<br>AAV x Epoch | $F_{1, 6}$ = 4.40<br>$F_{1, 6}$ = 5.83<br>$F_{1, 6}$ = 9.71 | < 0.081<br>< 0.052<br>< 0.021 | 0.42<br>0.49<br>0.62 |

Abbreviations: DV, dependent variable; $^E$Greenhouse-Geisser epsilon value
